# Novel Structural Features of Human Norovirus Capsid

**DOI:** 10.1101/528240

**Authors:** Jessica Devant, Götz Hofhaus, Grant S. Hansman

## Abstract

Human noroviruses are a major cause of gastroenteritis, yet there are still no vaccines or antivirals available. Nevertheless, a number of vaccine candidates that are currently in clinical trials are composed of norovirus virus-like particles (VLPs). These VLPs are recognized as morphologically and antigenically similar to norovirus virions. An X-ray crystal structure of the prototype (GI.1) VLPs showed that the norovirus capsid has a T=3 icosahedral symmetry and is composed of 180 copies of the major capsid protein (VP1) that folds into three quasi-equivalent subunits (A, B, and C). In this study, we determined the cryo-EM structure of VLPs for two GII.4 noroviruses that were detected in 1974 and 2012. We showed that these VLPs had a T=4 symmetry and were composed of 240 copies of VP1. The VP1 on the T=4 VLPs adapted four quasi-equivalent subunits (termed A, B, C, and D), which formed two distinct dimers (A/B and C/D). We found that the T=4 protruding domain was elevated ~21 Å off the capsid shell, which was ~7 Å more than the previously determined for the T=3 GII.10 norovirus. Another interesting feature of the T=4 VLPs was a small cavity and flap-like structure located at the twofold axis. This structural feature was associated with the shell domain (D subunit) and disrupted the contiguous shell. Altogether, we showed that the T=4 VLPs had a number of structural similarities and differences with other noroviruses, but how these structural changes associate with norovirus virions could be important for vaccine studies.

**IMPORTANCE:** The discovery that the GII.4 VLPs (identified in 1974 and 2012, termed CHDC-1974 and NSW-2012, respectively) have a T=4 symmetry is of major significance, since the NSW-2012 is clinically important and previous structural and biochemical studies assumed noroviruses have a T=3 symmetry and are composed of 180 copies of VP1. More importantly, NSW-2012 norovirus shared 96% amino acid identity with a GII.4 vaccine candidate and our data suggests that this vaccine might also have a T=4 symmetry. Although it is not clear if the T=4 VLPs were an artifact of the insect cell expression system, the T=4 VLP vaccines might not recognize equivalent epitopes on T=3 virions, which will be important for future neutralization studies. Finally, further studies with other norovirus genotypes and virions are clearly needed in order to determine the level of this structural diversity.

## INTRODUCTION

Human noroviruses are members of the *Caliciviridae* family and are a leading cause of outbreaks of acute gastroenteritis. The virus has a positive sense, single stranded RNA genome of ~7.7 kbp. The genome is organized into three open reading frames (ORFs), where ORF1 encodes nonstructural proteins and ORF2 and ORF3 encode a major structural protein (termed VP1) and a minor structural protein (termed VP2), respectively. Noroviruses are genetically diverse and based on VP1 sequences there are seven genogroups (GI-GVII), where GI, GII, and GIV cause infections in humans (1, 2). The GI and GII are further subdivided into numerous genotypes, with GII genotype 4 (GII.4) recognized as the most prevalent and clinically important (3, 4).

Development of an effective norovirus vaccine or antiviral has been hampered by the lack of a suitable cell culture system or small animal model. Moreover, the extensive genetic and antigenic diversity likely hinders vaccine development. Nevertheless, several vaccine candidates have progressed to phase I and II human clinical trials. Most vaccines were composed of norovirus GII.4 or GI.1 virus-like particles (VLPs), which can be produced by expressing VP1 in insect cells. These vaccines were well tolerated, highly immunogenic, and appeared to be safe, since they did not comprise of live or attenuated virus. However, limitations of the current vaccine formulations included mild norovirus like-symptoms, restricted long-term immunity, and limited cross-protection (5, 6).

Structural analysis of GI.1 VLPs reveals that VP1 is separated into two distinct domains: a shell domain (S domain) that encloses the RNA and a protruding domain (P domain) that binds to co-factors, such as histo-blood group antigens (HGBAs) and bile acids (7–9). A hinge region, which is typically composed of 10-14 amino acids, also connects the S and P domains. The P domain has a β-barrel fold that is structurally conserved in the *Caliciviridae* family. Dimerization of the P domains forms arch shaped protrusions that can be seen using electron microscopy. The P domain is further subdivided into P1 and P2 subdomain, where P2 subdomain appears to be an insertion in the P1 subdomain and is the most variable region on the capsid (7).

Structural studies have shown that caliciviruses have a common overall organization of T=3 icosahedral symmetry and are comprised of 180 copies of VP1 (7, 10–13). On the virus particles, the VP1 forms three quasi-equivalent subunits, termed A, B, and C (7). The norovirus A and B subunits assemble into 60 dimers (termed A/B) at the quasi twofold axis, whereas the C subunit assembles into 30 C/C dimers that are located at the strict icosahedral twofold axis. For the GI.1 VLPs, the A/B dimers have a bent S domain conformation, whereas the C/C dimers have flat S domain conformation (7). The conformational differences within these dimers likely facilitates the curvature of the virus particle to form a closed shell, which is commonly seen in other T=3 icosahedral viruses (14). Interestingly, smaller norovirus VLPs (~25 nm in diameter) that are assumed to have a T=1 icosahedral symmetry were also reported (15, 16); however the structure of these smaller VLPs has not yet been determined.

In this study, we determined the cryo-EM structure of VLPs for GII.4 variants that were identified in 1974 and 2012, termed CHDC-1974 and NSW-2012, respectively (17, 18). We showed that these VLPs had a T=4 icosahedral symmetry and were composed of 240 copies of VP1. In order to form the T=4 symmetry, VP1 adapted four quasi-equivalent subunits, termed A, B, C, and D, which subsequently gave rise to two distinct dimers, termed A/B and C/D. These VLPs consisted of 60 A/B dimers and 60 C/D dimers, where at the icosahedral 2-fold axis, B, C, and D subunits were alternating, while the A subunit was located at the fivefold axis. Altogether, our findings showed that the GII.4 VLPs had structural modifications that might have important implications for vaccine design.

## MATERIALS AND METHODS

### VLP preparation

The NSW-2012 and CHDC-1974 VLPs (Genebank accession numbers JX459908 and ACT76142, respectively) were expressed in a baculovirus system as previously described (19–21). Briefly, the bacmid containing the recombinant VP1 gene was transfected in Sf9 insect cells. After incubation for five days, the culture medium was centrifuged for 10 min at 3,000 rpm at 4°C. The recovered baculovirus was subsequently used to infect Hi5 insect cells. After five days post infection, the culture medium was centrifuged for 10 min at 3,000 rpm at 4°C and then 1 h at 6,500 rpm at 4°C. The VLPs in the supernatant were concentrated by ultracentrifugation at 35,000 rpm for 2 h at 4°C and then further purified using CsCl equilibrium gradient ultracentrifugation at 35,000 rpm for 18 h at 4°C. To remove the CsCl, the VLPs were pelleted for 2 h at 40,000 rpm at 4°C and subsequently resuspended in PBS (pH 7.4).

### Negative stain electron microscopy

The integrity of the VLPs was confirmed by negative stain electron microscopy (EM). GII.4 virions from stool were also examined using EM (prepared as above expect for the CsCl equilibrium gradient ultracentrifugation step). Briefly, the VLPs were diluted 1:30 in distilled water and applied to EM grids, whereas the stool sample was diluted to 10% with PBS, applied to EM grid, washed with water, and then fixed with 4% glutaraldehyde. The grids were washed with distilled water, stained with 0.75% uranyl acetate, and the excess uranyl acetate removed with filter paper. EM images were acquired on a Zeiss 910 electron microscope at 50,000× magnification.

### Cryo-EM data sample preparation and data collection

UNSW-2012 and CHDC-1974 VLPs (3 μl) were applied on freshly glow discharged Quantifoil holey carbon support films (R1.2/1.3; Quantifoil) and blotted for 18 seconds at 100% humidity at 10°C before been plunged in liquid ethane using an FEI Mark IV Vitrobot (Thermo Fischer Scientific). Vitrified specimens were imaged on a Titan Krios operated at 300 keV. NSW-2012 micrographs were acquired with a K2 direct electron detector at 64,000× magnification, corresponding to a pixel size of 2.27 Å/px, while CHDC-1974 micrographs were collected using a K3 camera with Latitude S software (Gatan) at 64,000× magnification, corresponding to a pixel size of 1.375 Å/px.

### Cryo EM data processing

Images were processed with Relion 2.1 software for NSW-2012 and cryosparc software for CHDC-1974 (22, 23). Initially, the movies containing 16 frames for NSW-2012 and 40 frames for CHDC-1974 were motion corrected using motioncor2 software (24) and contrast transfer function (CTF) was performed using ctffind 4.1 software (25). An initial set of 1,000 particles was manually picked for 2D classification to produce averages suitable as references for automated particle picking. The autopicked particles were sorted in a 2D classification step and the best particles were used for calculation of an initial starting model, followed by 3D classification. A subset of particles that generated the highest resolution was selected for further refinement. The 3D refinement and post-processing of NSW-2012 from 10,548 particles produced a final map at 7.3-Å resolution with I2 symmetry imposed (0.143 FSC cutoff). For CHDC-1974, a subset of 42,485 particles for refinement revealed a map of 6.1-Å resolution using the 0.143 FSC cutoff. Cryo-EM VLP structures for CHDC-1974 (accession number: EMD-4549) and NSW-2012 (EMD- 4550) were deposited at EMDB.

### Fitting of the X-ray structures into the density maps

Crystal structures of NSW-2012 P domain (PDB ID: 4OOS) and CHDC-1974 P domain (5IYN) were fitted into the respective densities using the “fit in map” command in the UCSF Chimera software (26). Since a high-resolution GII.4 shell domain was unavailable, the GI.1 Norwalk virus S domain was extracted from the X-ray crystal structure (1IHM) and fitted into the GII.4 cryo-EM density using UCSF Chimera software.

## RESULTS AND DISCUSSION

The purpose of this study was to analyze the GII.4 VLP architecture of NSW-2012 and CHDC-1974 and then relate the findings with known GI.1 and GII.10 VLP structures. The NSW-2012 VP1 sequence had a single amino acid insertion at position ~394 (NSW-2012 numbering) compared to CHDC-1974 (Fig. 1). Overall, NSW-2012 and CHDC-1974 shared 89% amino acid identity. Most (45 of 54) amino acid substitutions were located in the P domain. Negative stain EM images revealed that these the VLPs exhibited characteristic norovirus morphology (Fig. 2). However, the diameter of these VLP images was measured to be ~52 nm, which suggested that the GII.4 VLPs were larger than GII.10 and GI.1 VLPs that had diameters of ~43 nm and 38 nm, respectively.

**Figure 1.**
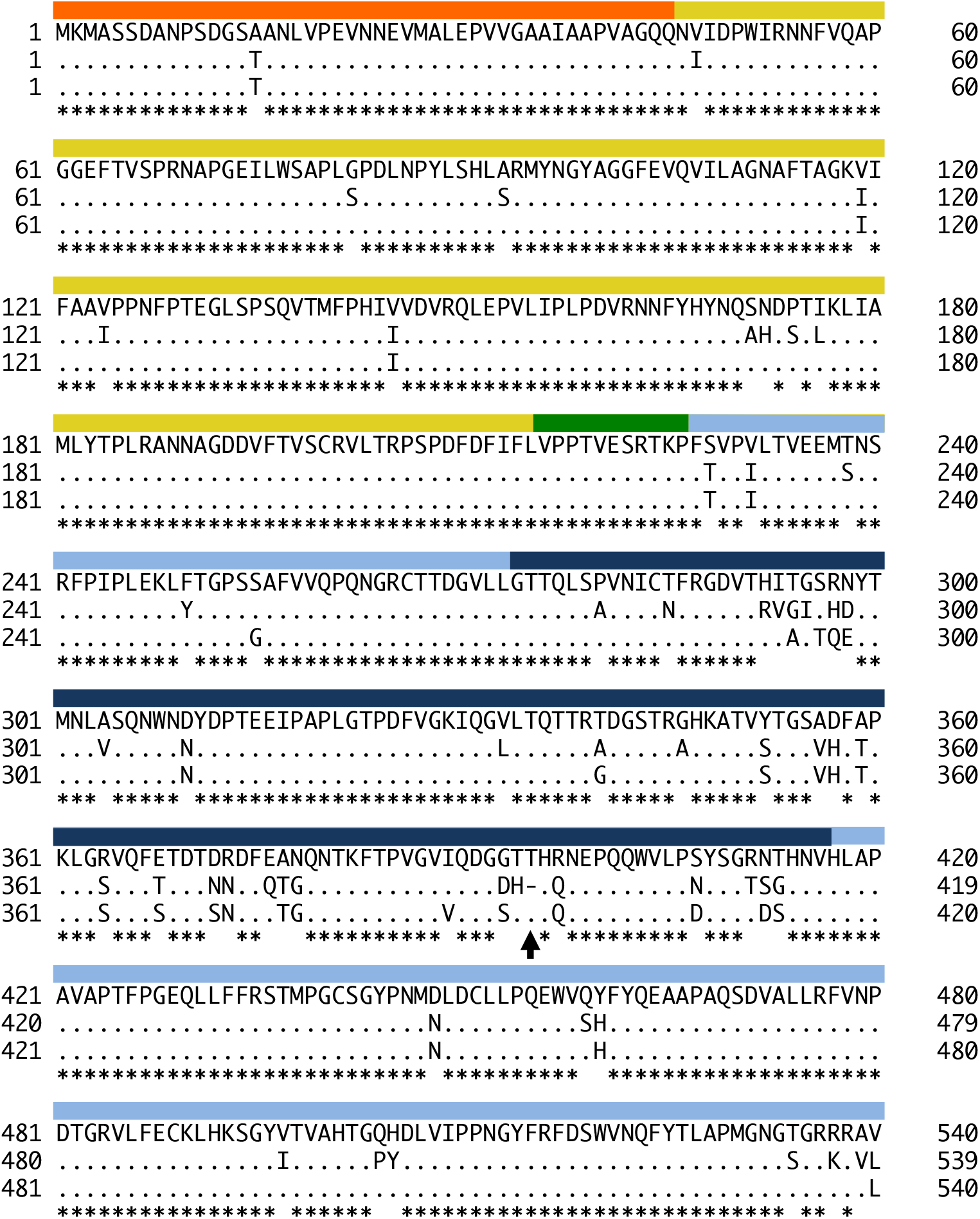
Amino acid sequence alignment of GII.4 VP1. NSW-2012 (JX459908), CHDC-1974 (ACT76142), and GII.4c (31) VP1 amino acid sequences were aligned using ClusterX. The S domain (orange), hinge region (green), P1 subdomain (light blue), and P2 subdomain (navy) were labeled accordingly. Compared to CHDC-1974, NSW-2012 and GII.4c VP1 had a single amino acid insertion (arrow) at position 394 (NSW-2012 numbering). The S domain and hinge region were mainly conserved, whereas most amino acid substitutions were located in the P2 subdomain.

**Figure 2.**
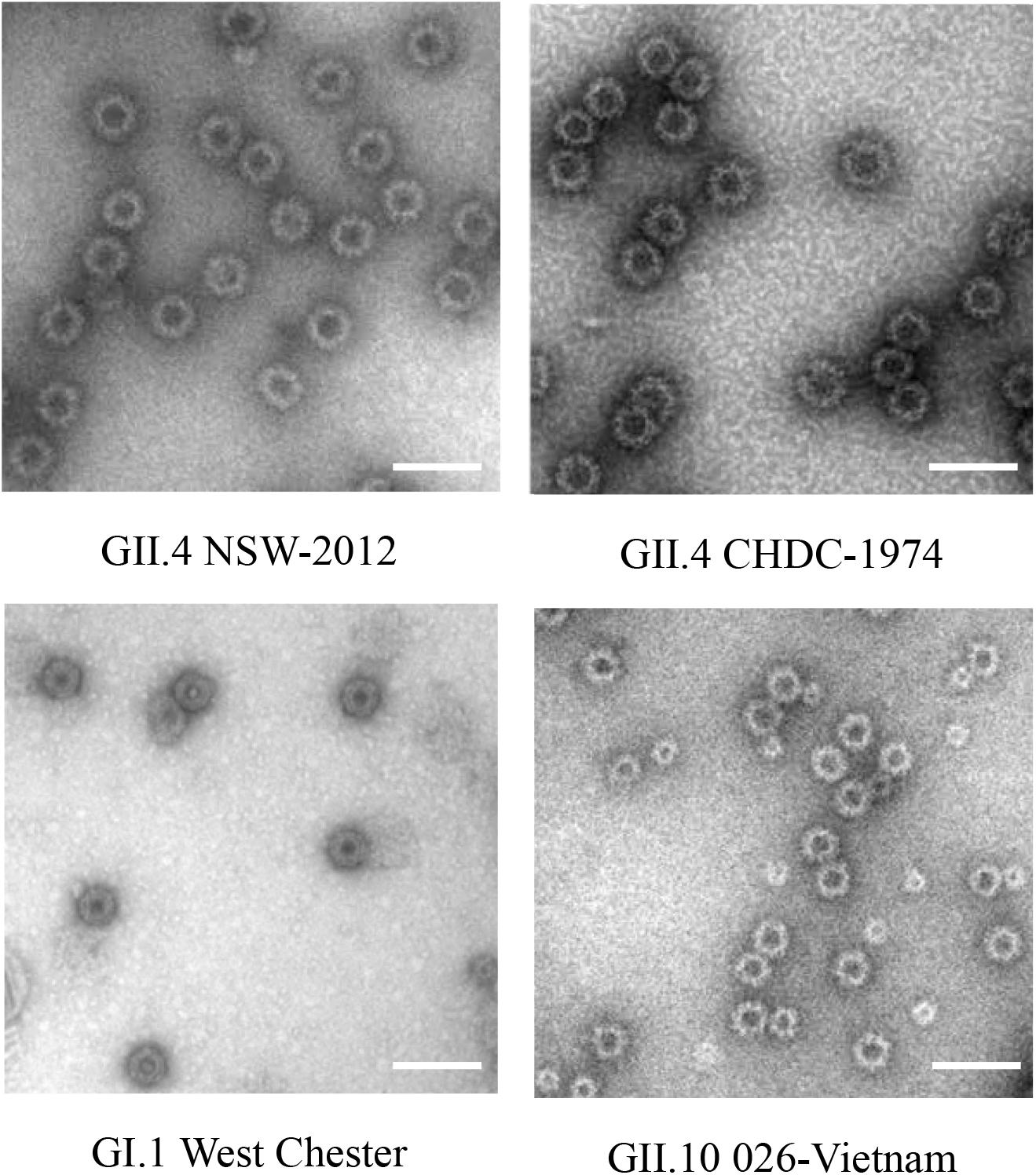
EM images and hydrodynamic diameters of NSW-2012 and CHDC- 1974 GII.4 VLPs. Negative stain EM images of the GII.4 VLPs show the characteristic norovirus morphology (50,000× magnification). The GI.1 West Chester (AY502016.1) and GII.10 Vietnam026 (AF504671) VLPs are shown as reference (27, 34). The bar represents 100 nm.

### Cryo-EM structure of GII.4 NSW-2012 VLPs

The structure of the NSW-2012 VLPs was determined using cryo-EM. The VLPs were mono-dispersed in vitreous ice and appeared homogenous in size (Fig. 3A). From the 364 images, 10,548 particles were used for structural reconstruction and refined to 7.3 Å resolution (Fig. 3B).

**Figure 3.**
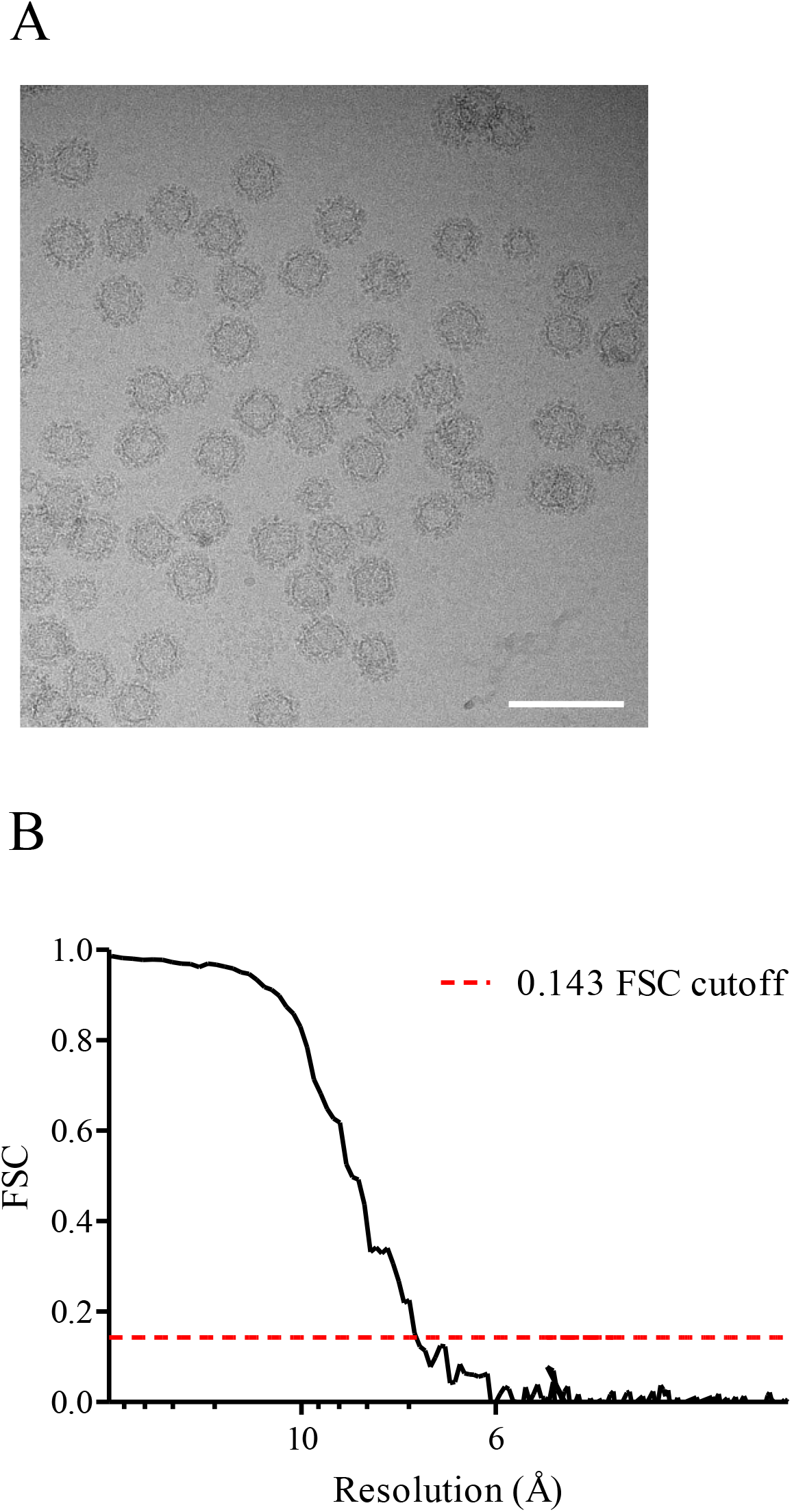
NSW-2012 cryo-EM data processing. (A) A representative cryo-EM micrograph of NSW-2012 VLPs at 64,000× magnification. The scale bar represents 100 nm. (B) Gold standard Fourier shell correlation (FSC) plot of the icosahedral reconstruction of NSW-2012 indicates a resolution of 7.3 Å.

Unexpectedly, NSW-2012 VLPs were discovered to have a T=4 icosahedral symmetry (Fig. 4). Our data revealed that these VLPs were composed of 240 copies of VP1, rather than the 180 VP1 copies in GI.1 and GII.10 VLPs. The inner diameter of NSW-2012 shell was measured as 32 nm, while the outer capsid diameter was 50 nm. Interestingly, this large diameter of the T=4 VLPs corresponded to an inner shell volume of 17,157 nm^3^, which was ~2.6 times the volume of the GII.10 VLPs that had an inner diameter of 23 nm (13).

**Figure 4.**
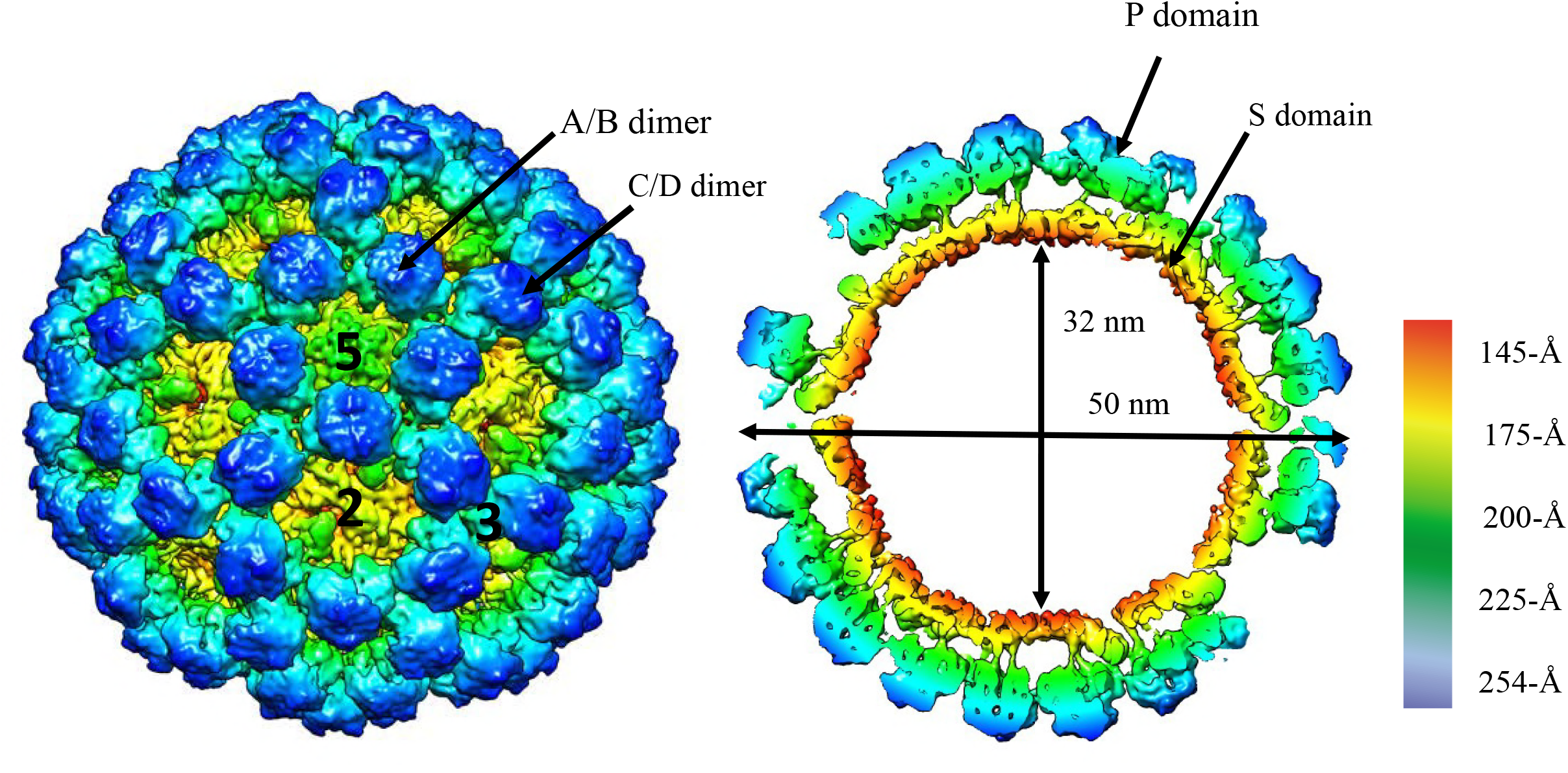
Cryo-EM reconstruction structure of NSW-2012 VLPs. The left side shows NSW-2012 VLPs have a T=4 icosahedral symmetry (symmetry axis labeled 2, 3, and 5). These VLPs were composed of 240 copies of VP1. The VP1 adapted four quasiequivalent conformations (A, B, C, and D) that gave rise to two distinct dimers (A/B and C/D). At the icosahedral twofold axis, the B, C, and D subunits were alternating, while the A subunit is positioned at the fivefold axis. The right side shows a cutaway section of these VLPs and indicates that the inner and outer diameters are 32 nm and 50 nm, respectively. The P domains are elevated ~21 Å off the S domain.

Another interesting structural feature of these NSW-2012 VLPs was a small cavity and flap-like structure on the contiguous shell (Fig. 5). This feature was associated with the S domain and found on opposing sides at the twofold axis. Interestingly, NSW-2012 VLPs were capable of binding HBGAs and norovirus-specific antibodies (27, 28), which indicated that despite the cavity and flap-like structure these VLPs were biologically functional.

**Figure 5.**
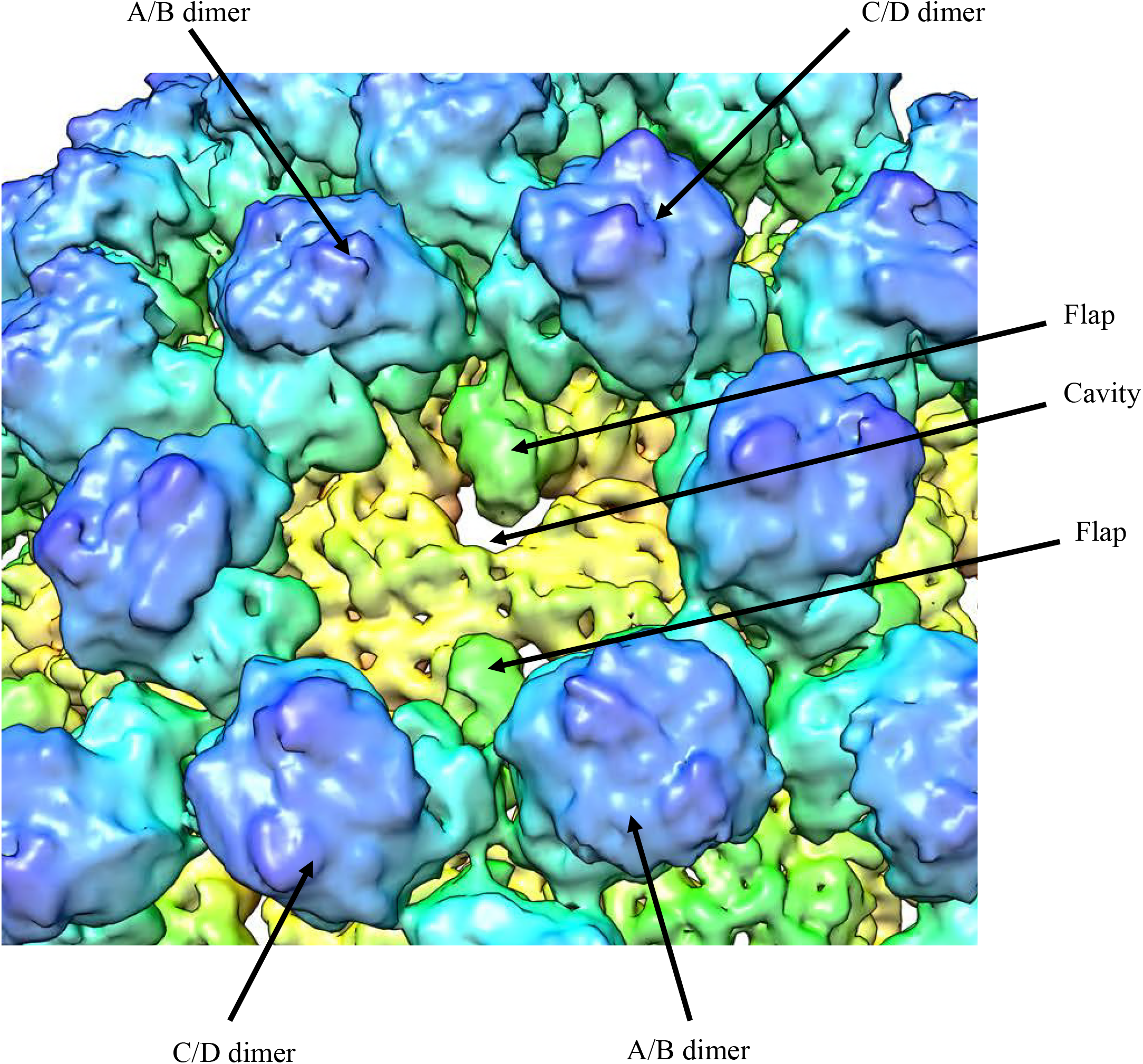
NSW-2012 T=4 VLPs shows several new structural features. The cavity and flap-like structures are observed at the twofold axis and are found on opposing sides. The cavity and flap-like structures are associated with the S domain on the D subunit.

Structural analysis of the T=4 VLPs indicated that VP1 adapted four quasi-equivalent subunits, termed A, B, C, and D. Subsequently, these four subunits gave rise to two distinct dimers, termed A/B and C/D (Figs. 4 and 6). At the icosahedral twofold axis, B, C, and D subunits were alternating, while the A subunit was positioned at the fivefold axis (Fig. 4). Of major importance, the T=4 VLPs comprised of 60 A/B dimers and 60 C/D dimers, which was distinct from the T=3 GI.1 and GII.10 VLPs that assembled with 60 A/B and 30 C/C dimers. We also observed that the T=4 A/B and C/D dimers had a bent conformation at the bottom of the S domain, which was in contrast to the GI.1 VLPs that consisted of both bent (A/B) and flat (C/C) dimers (7). Likely, the bent A/B and C/D dimers facilitated the necessary curvature to form particles with a T=4 symmetry.

**Figure 6.**
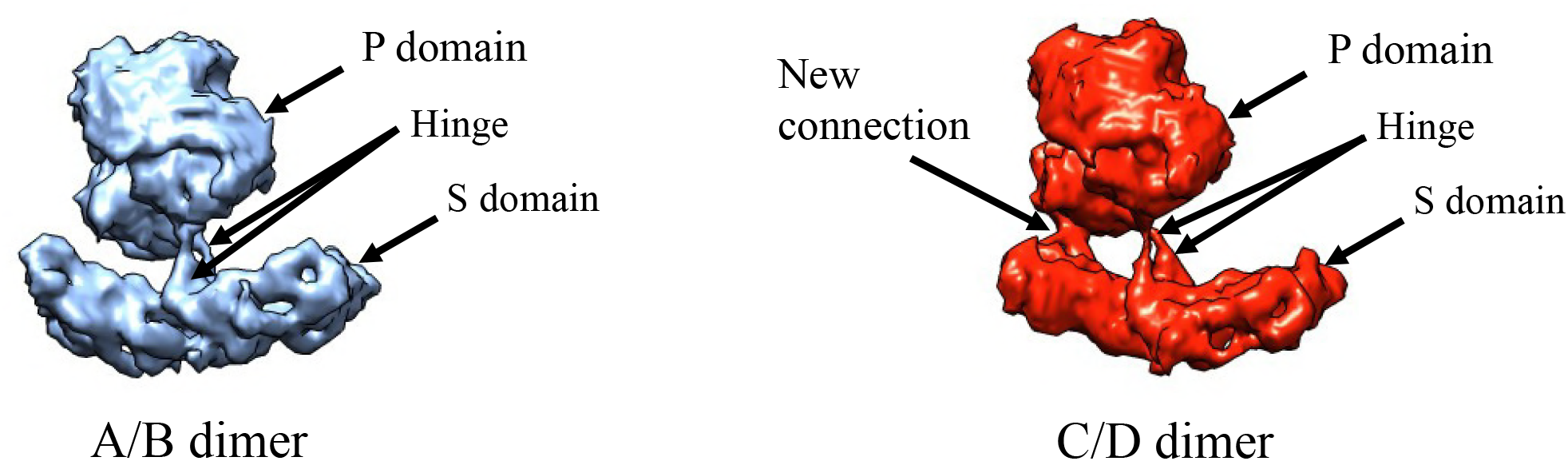
NSW-2012 T=4 VLPs are formed with 60 A/B and 60 C/D VP1 dimers. The A/B and C/D dimers show an equivalent bent confirmation on the bottom of the S domain. An additional connection was observed between the D subunit of the S and P domain.

In order to better comprehend how VP1 assembled into the T=4 VLPs, the X-ray crystal structure of NSW-2012 P domain (4OOS) and GI.1 S domain (1IHM) were fitted into the VLP density map. We found that the NSW-2012 P domain dimer could be unambiguously positioned into the VLP structure with cross correlation coefficient of 0.96 (Fig. 7A). This result indicated that the P domain dimers on the T=4 VLPs had not undergone any major structural modifications. In the case of the S domain, the GI.1 S domain needed to be manually positioned into the density map. The GI.1 S domain fitted well into to A/B dimer and the C subunit, while the D subunit needed to be further repositioned (Fig. 7B). This additional fitting in the D subunit was necessary in order to occupy the elevated density of the cavity and flap-like regions.

**Figure 7.**
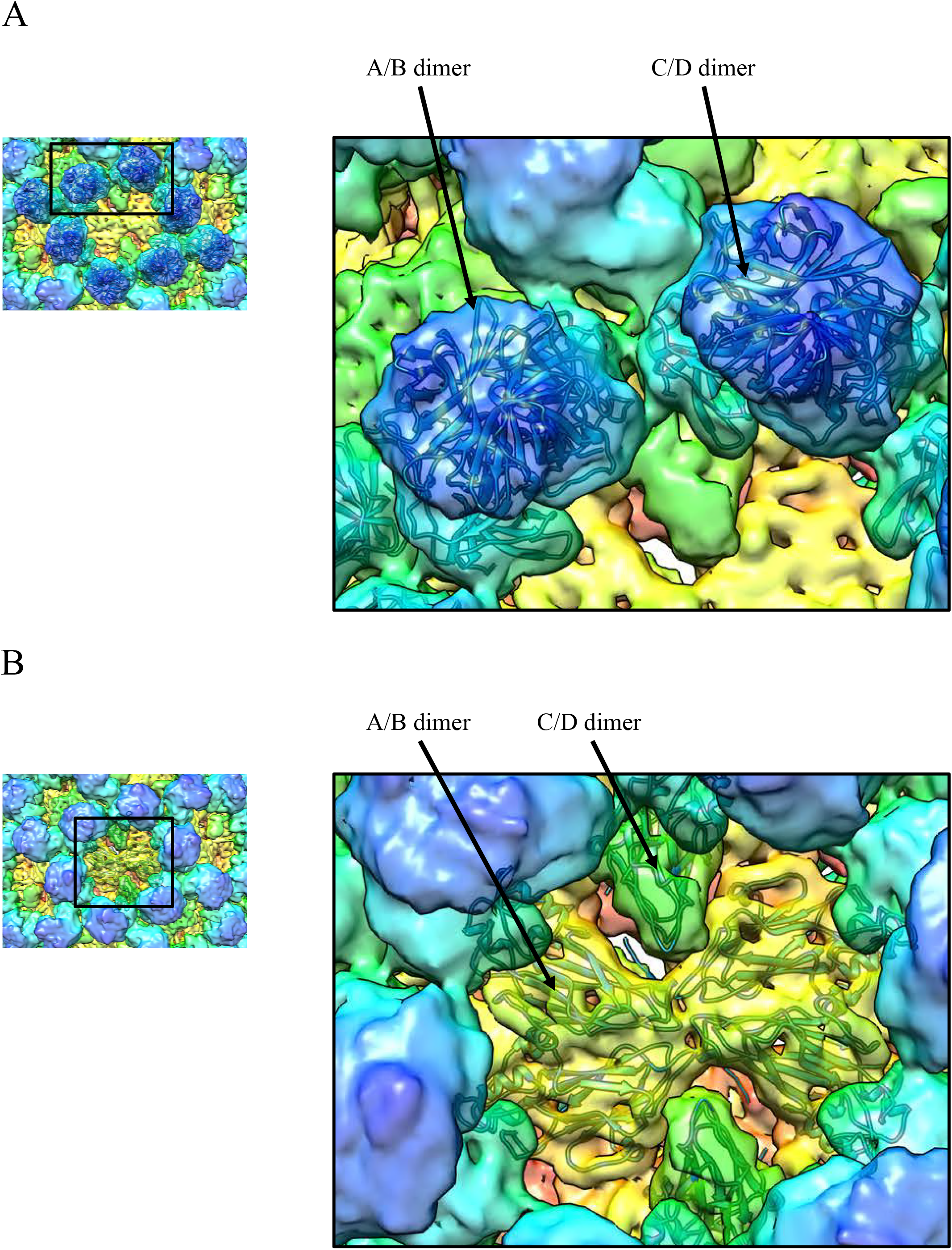
The X-ray crystal structures of NSW-2012 P domain and GI.1 S domain were fitted into the VLP density map. (A) The X-ray crystal structure of NSW-2012 P domain (4OOS, cartoon) could be fitted into the A/B and C/D P domain dimers, indicating little conformational change. (B) The X-ray crystal structure of the GI.1 S domain (1IHM, cartoon) fitted into the A/B and C/D S domain dimers. However, the cavity and flap-like structures on the D subunit suggests a large conformational change compared to the typical T=3 particles.

Unfortunately, it was problematic to fit the hinge region, since the hinge region on the X-ray crystal structure of NSW-2012 P domain was excluded from the expression construct and the hinge region on the GI.1 VLPs was flattened (7, 13). Interestingly, an additional density was also observed between the S domain and the C-terminus of the P domain on the D subunit (Fig. 8). This connection appeared to stabilize the bent conformation of the C/D dimer and subsequently the T=4 VLPs. Indeed, the C- terminus of VP1 on the GI.1 VLPs and the GII.4 P domain were found to be flexible (7, 28). Interestingly, the C-terminus of VP1 was previously shown to be important for the size and stability of VLPs (29). Therefore, it is possible that GII.4 VP1 sequence has a remarkable ability to form T=4 VLPs, which could provide an additional advantage for this genotype.

**Figure 8.**
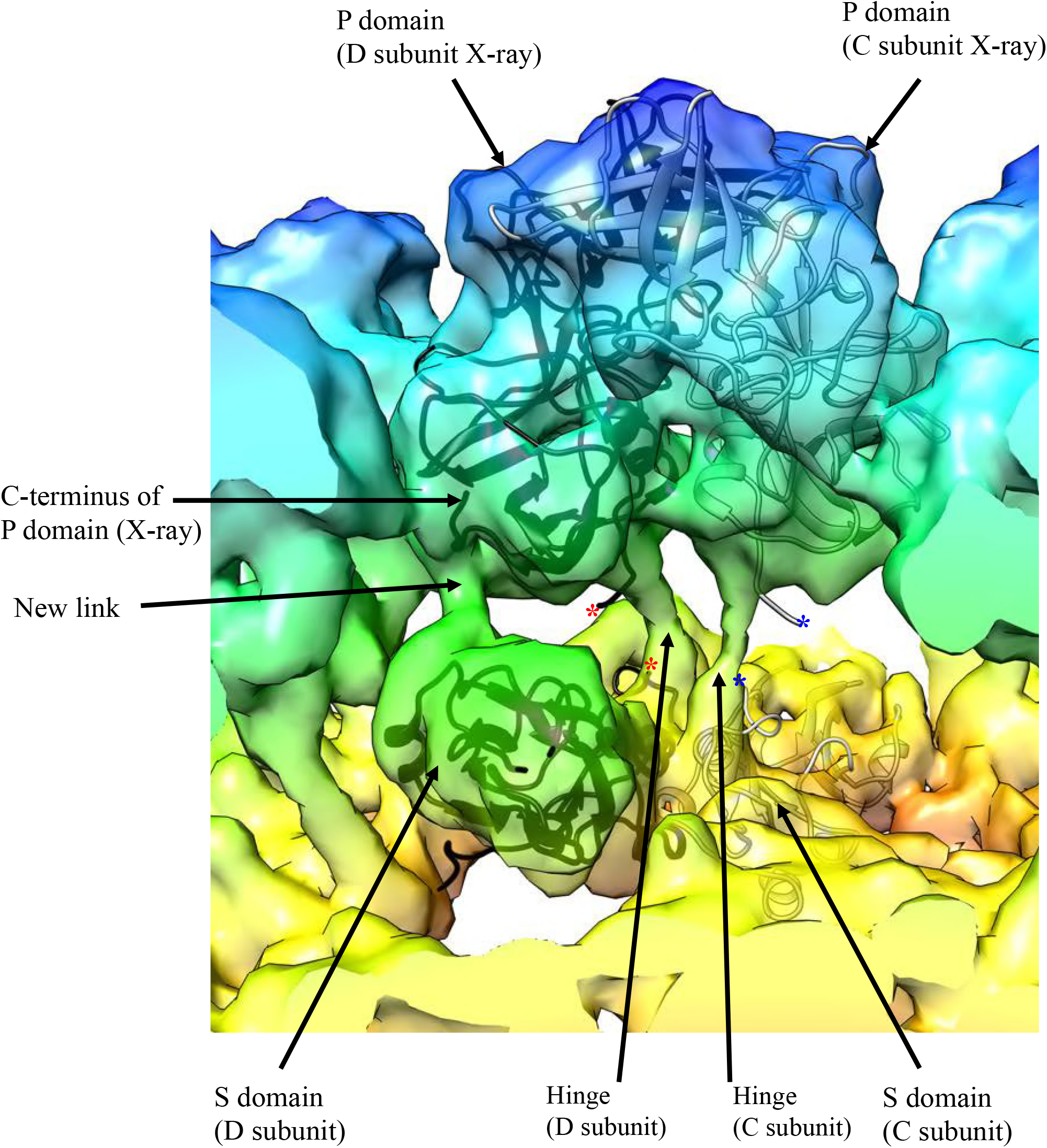
A close-up view of NSW-2012 C/D dimer. The fitted X-ray crystal structures of the GI.1 S domain (cartoon) and the GII.4 P domain (cartoon) into the cryo-EM map shows the how the hinge region connects the S and P domains. Also, the new connection between the S domain and the C-terminus of the P domain is shown. The asterisk represents the missing hinge region on the X-ray crystal structures that connects of the S and P domains for the C subunit (blue) and D subunit (red).

Another interesting feature that we observed with the T=4 VLPs was the raised P domains (Figs. 4 and 8). We found that the T=4 P domain was elevated ~21 Å off the shell, which was higher than the P domains on the GII.10 VLPs, which were raised ~14 Å (13). The hinge region in NSW-2012 and GII.10 (13) were both ~10 amino acids and mainly conserved (30). This result suggested that the raised P domains might be a structural feature of GII noroviruses, since the P domains on the GI.1 VLPs were essentially resting on the shell (7).

### Cryo-EM structure of GII.4 CHDC-1974 VLPs

Following these results, we proceeded to determine the cryo-EM structure of the CHDC-1974 VLPs. The VLPs were mostly mono-dispersed and homogenous in size (Fig. 9A). From 591 images, 42,485 particles were used for the structural reconstruction that led to a final resolution of 6.1 Å (Fig. 9B). Remarkably, the CHDC-1974 VLPs also had a T=4 symmetry (Fig. 10). In fact, the CHDC-1974 VLP structure closely resembled the structural features of NSW-2012 VLPs.

**Figure 9.**
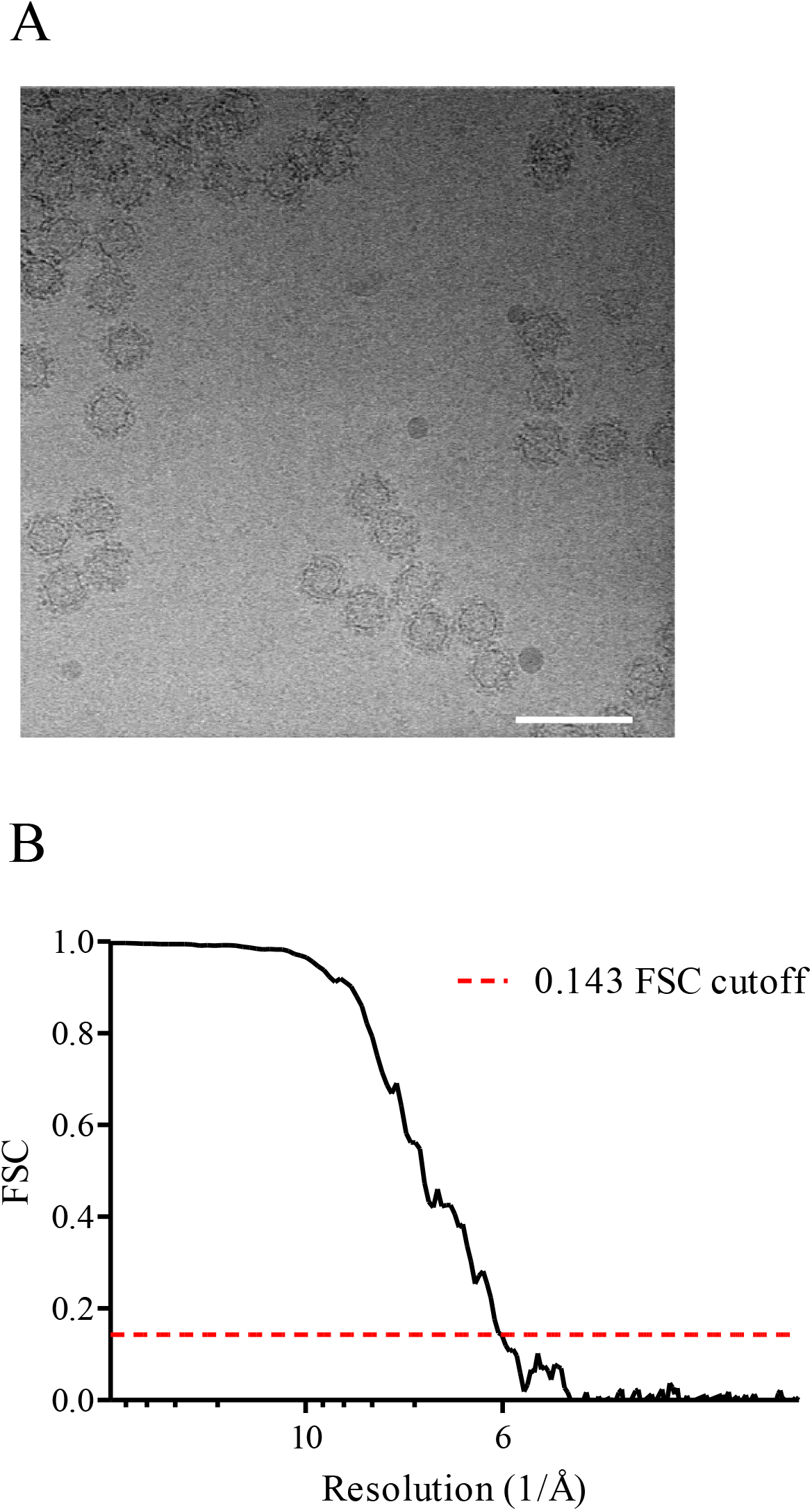
CHDC-1974 cryo-EM data processing. (A) A representative cryo-EM micrograph of CHDC-1974 VLPs at 64,000× magnification. The scale bar represents 100 nm. (B) FSC plot of the icosahedral reconstruction of CHDC-1974 indicates a resolution of 6.1 Å at 0.143 cutoff.

**Figure 10.**
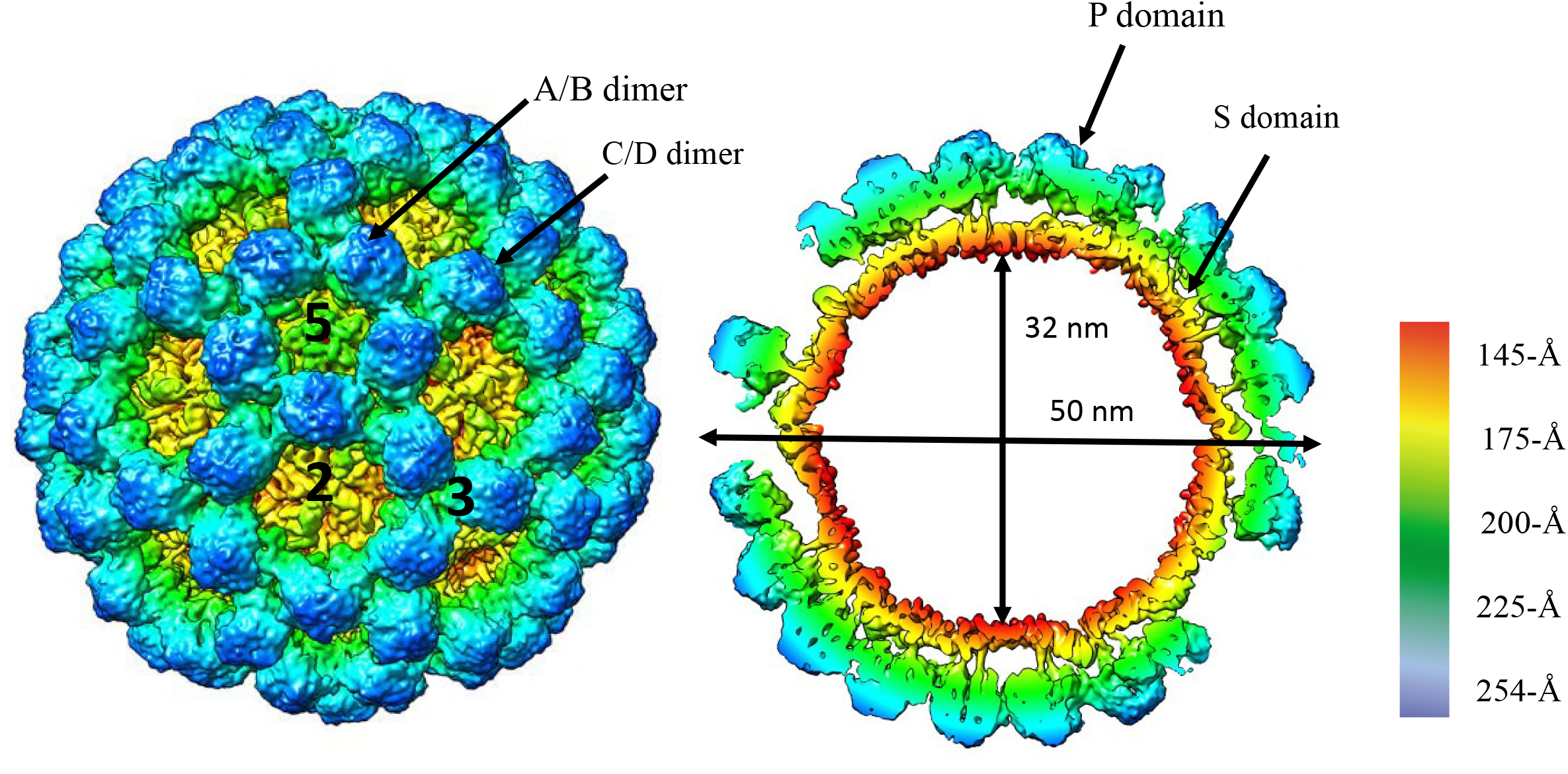
Cryo-EM structure and analysis of CHDC-1974 VLPs. The image on the left side shows that CHDC-1974 VLPs has a T=4 icosahedral symmetry and was composed of 240 copies of VP1. The VP1 adapted four quasiequivalent conformations (A, B, C, and D) that gave rise to two distinct dimers (A/B and C/D). At the icosahedral twofold axes, the B, C, and D subunits were alternating, while the A subunit was located around the fivefold axes. The right side shows a cutaway section of these VLPs and indicates that the inner and outer diameters are 32 nm and 50 nm, respectively. The P domains are elevated ~21 Å off the S domain.

We found that CHDC-1974 VLPs were composed of 240 copies of VP1 that formed the equivalent subunits (A, B, C, and D) and A/B and C/D dimers. The inner diameter of the shell was 32 nm, whereas the outer diameter of the capsid was 50 nm. The comparable small cavity and flap-like structures were also present on the CHDC-1974 VLPs (Fig. 11). The CHDC-1974 A/B and C/D dimers showed a similar bend as NSW-2012 dimers, although slightly less pronounced (Fig. 12). The X-ray crystal structure of CHDC-1974 P domain (5IYN) was easily fitted into the CHDC-1974 VLP density map (Fig. 13A). The GI.1 S domain also fitted into the A, B, and C subunits, whereas the GI.1 S domain needed to be again repositioned to occupy the D subunit (Fig. 13B). Similar to NSW-2012 VLPs, an additional density was observed between the S and P domains on the D subunit (Fig. 14). Lastly, we found that CHDC-1974 P domain was also lifted off the shell by ~21 Å (Figs. 10 and 14).

**Figure 11.**
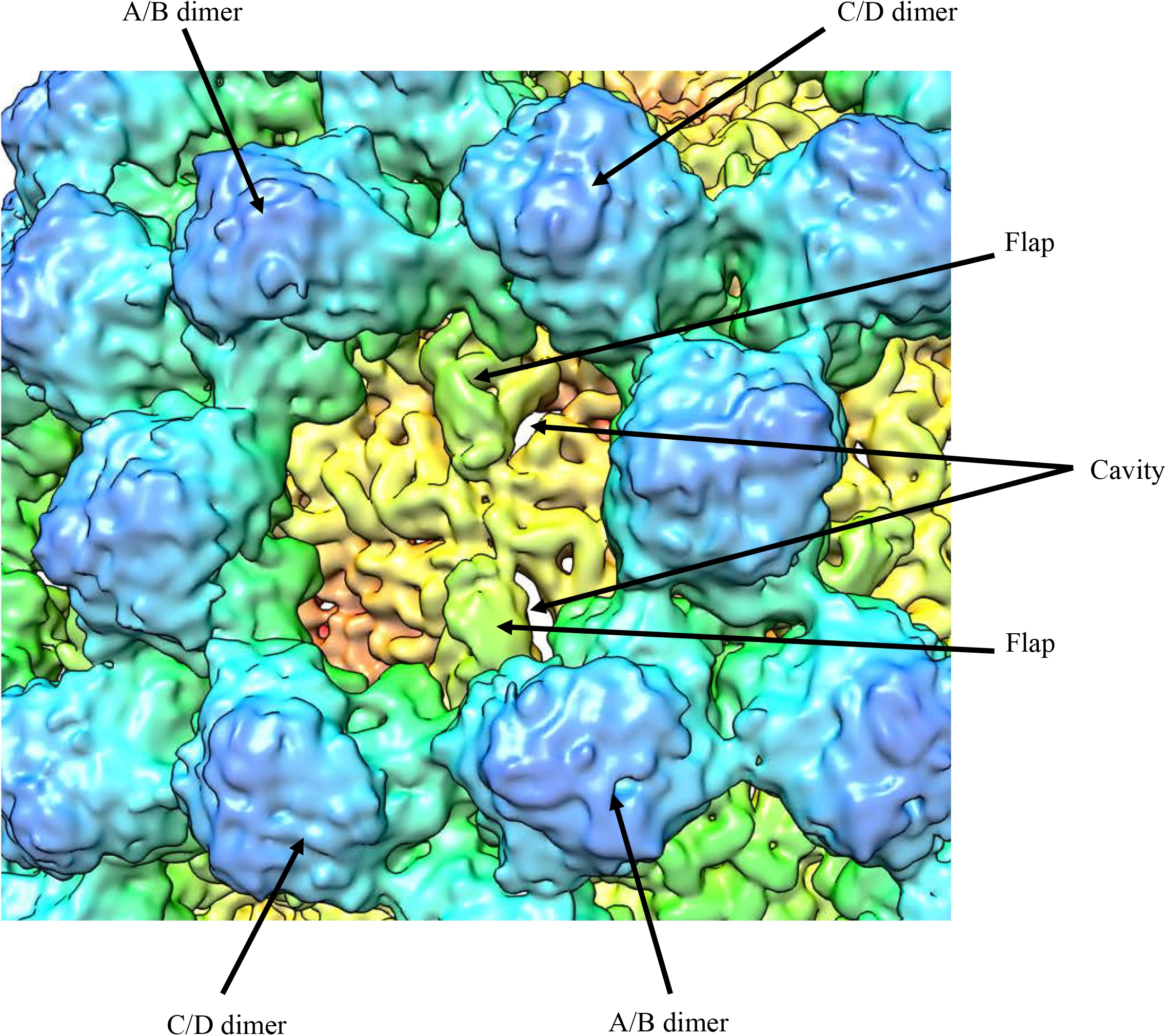
CHDC-1974 T=4 VLPs shows several new structural features. The cavity and flap-like structures are observed at the twofold axis and are found on opposing sides. The cavity and flap-like structures are associated with the S domain on the D subunit.

**Figure 12.**
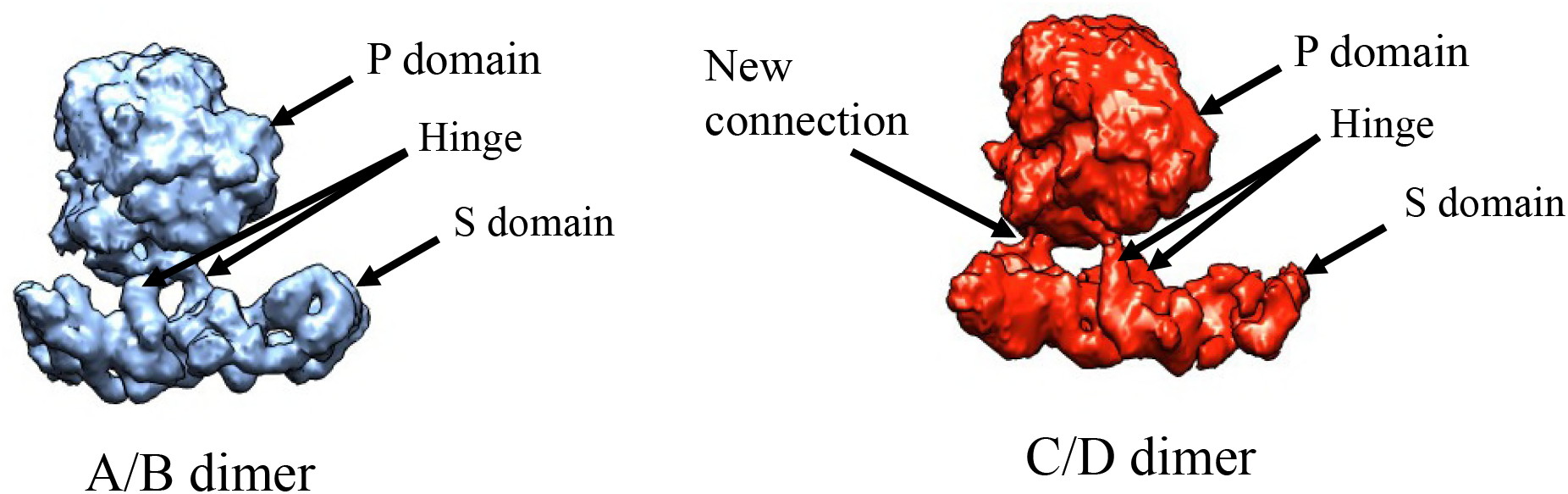
CHDC-1974 T=4 VLPs are formed with 60 A/B and 60 C/D VP1 dimers. The A/B and C/D dimers show an equivalent bent confirmation on the bottom of the S domain. An additional connection was observed between the D subunit of the S and P domain.

**Figure 13.**
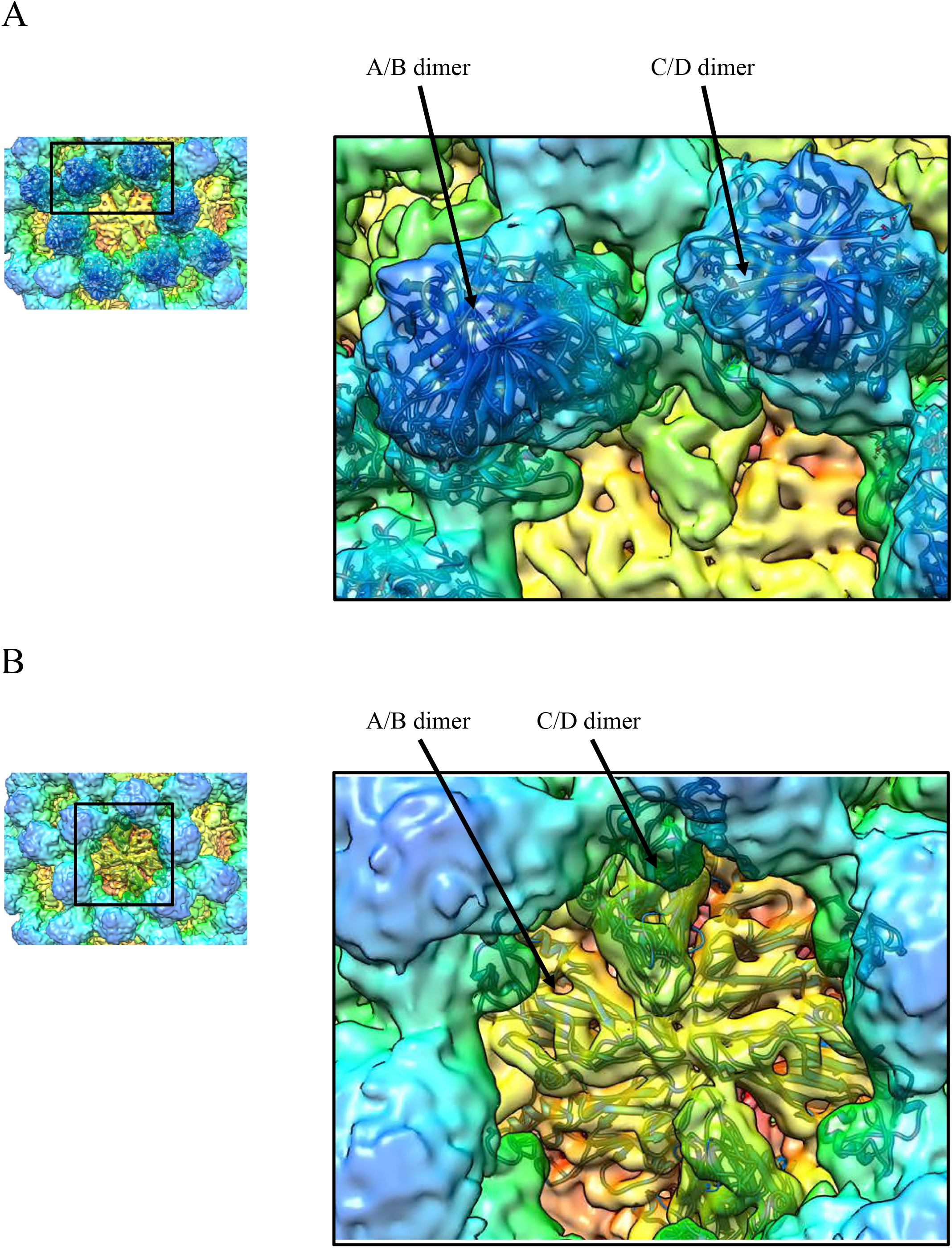
The X-ray crystal structures of CHDC-1974 P domain and GI.1 S domain were fitted into the VLP density map. (A) The X-ray crystal structure of NSW-2012 P domain (5IYN, cartoon) easily fitted into the A/B and C/D P domain dimers. (B) The X-ray crystal structure of the GI.1 S domain (1IHM, cartoon) fitted into the A/B and C/D S domain dimers. However, the cavity and flap-like structures on the D subunit suggests a large conformational change from typical T=3 particles.

**Figure 14.**
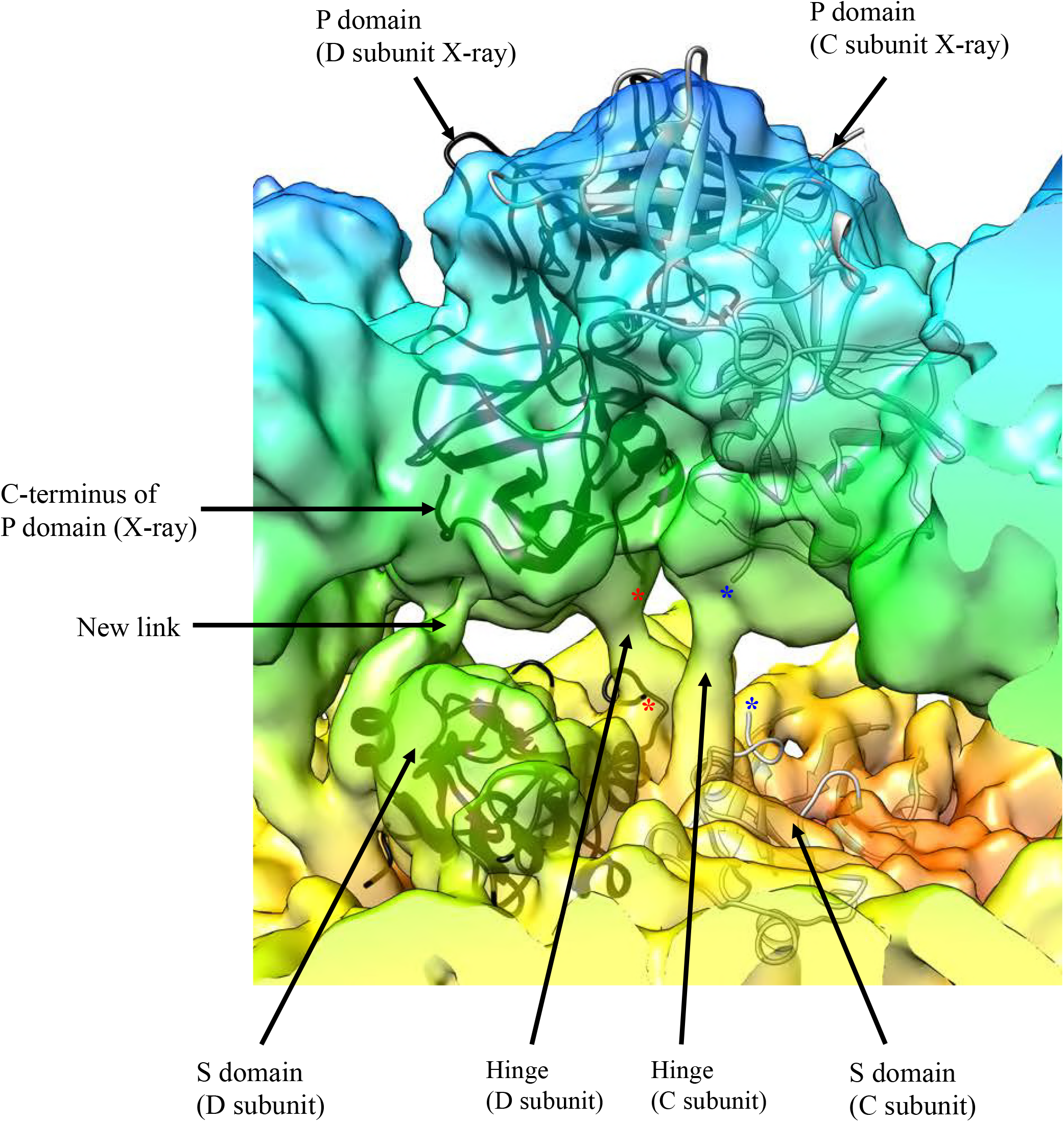
A close-up view of CHDC-1974 C/D dimer. The fitted X-ray crystal structures of the GI.1 S domain (cartoon) and the GII.4 P domain (cartoon) into the cryo-EM map shows the how the hinge region connects the S and P domains. Also, the new connection between the S domain and the C-terminus of the P domain is shown. The asterisk represents the missing hinge region on the X-ray crystal structures that connects of the S and P domains for the C subunit (blue) and D subunit (red).

Overall, these results showed that GII.4 VP1 sequences isolated over three decades apart remained structurally conserved. This could imply that other GII.4 VP1 sequences also form T=4 VLPs when expressed in insect cells, especially since these two sequences had only 89% amino acid identity. More importantly, the GII.4 noroviruses have dominated epidemics over the past decade and this structural feature could represent a selective advantage over other GII genotypes that have a T=3 capsid.

### Structural implications of the T=4 VLPs

Our new discovery of these T=4 GII.4 VLPs could have major implications for vaccine development. Negative stain EM images of the GII.4c VLPs that are currently tested in clinical trials showed the typical norovirus morphology (6, 31). However, the size determination and the structure are not available.

In general, studies have shown that norovirus-specific antibody titers were raised after vaccination with VLPs, but the levels of protection were not strongly improved compared to placebo groups (32). It is tempting to speculate that the GII.4c VLPs might also form T=4 particles, since the GII.4c and NSW-2012 shared 94% amino acid identity; most (28 of 31) of the substitutions were located in the P domain; and the hinge region was identical (Fig. 1). Therefore, this might suggest that the host could produces neutralizing antibodies against epitopes on the T=4 VLPs that were not as accessible on T=3 virions and that efficacy is difficult. Indeed, we have identified several Nanobodies that bind to occluded regions on the T=4 GII.4 VLPs (30, 33).

In order to validate if the GII.4 virions actually assemble into T=3 a cryo-EM structure would be valuable, however norovirus virions are challenging to prepare in large quantities. Nevertheless, when we compare EM images of GII.4 virions with these T=4 VLPs we found that the virions were smaller (Fig. 15). Therefore, our preliminary results suggest that the GII.4 VLPs and virions were different in size and likely other structural characteristics.

**Figure 15.**
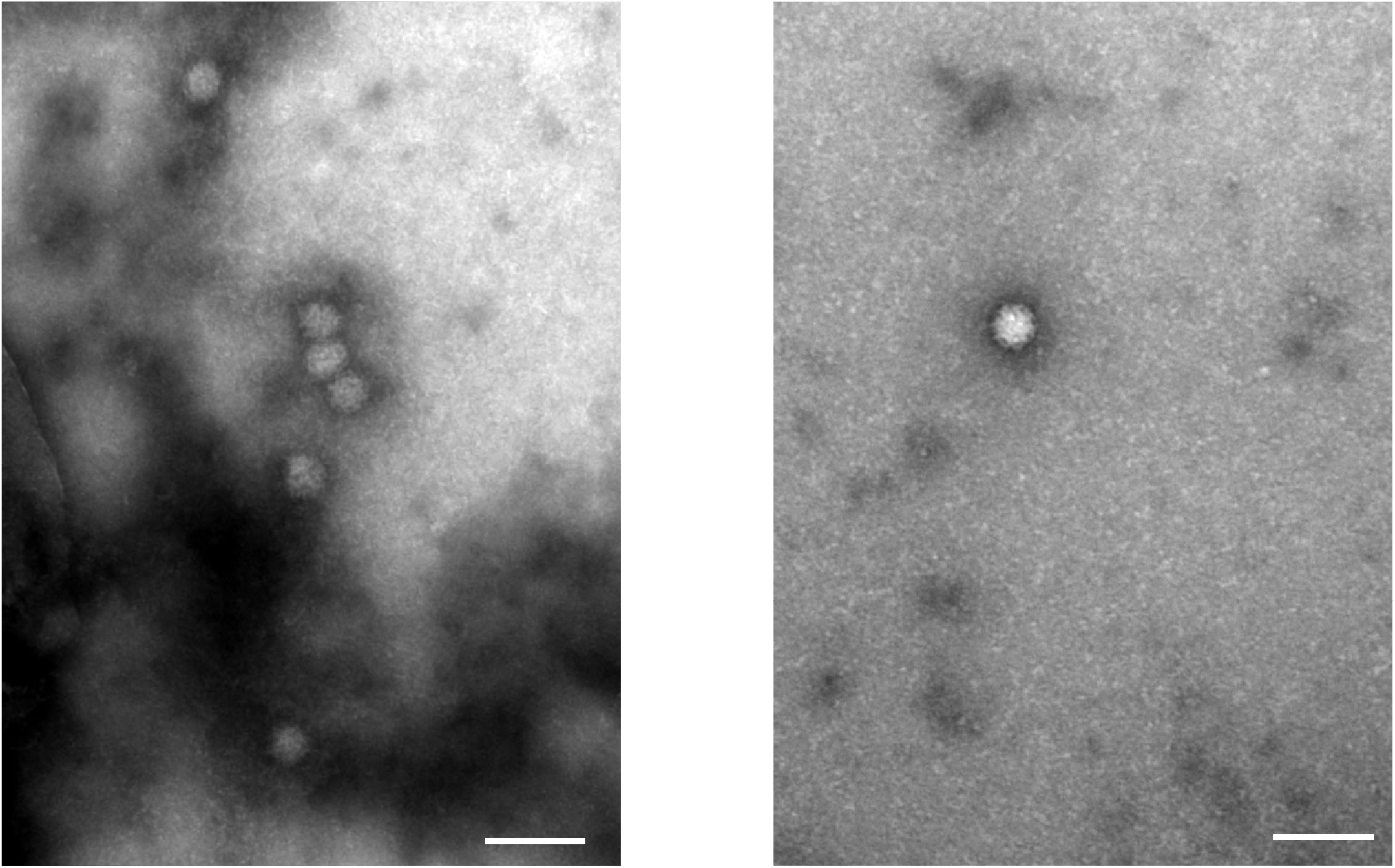
EM images of GII.4 virions. Negative stain EM images of GII.4 virions show that the virions exhibit a smaller diameter than GII.4 VLPs expressed in insect cells.

### Summary

At least two important outcomes from this new discovery of these GII.4 T=4 VLPs will be acknowledged. Firstly, the GII.4 VLPs might influence results from previous studies that assumed norovirus VLPs were morphologically similar to virions. For example, the molecular weight of the T=3 and T=4 VLPs would be ~10.5 mDa (180 × VP1) and ~14 mDa (240 × VP1), respectively. This mass difference would affect results in mass spectrometry, Biacore, and isothermal titration calorimetry. Secondly, the VLP vaccines that could be composed of T=4 particles might produce complicating immune responses. For example, the cavity and flap-like structures on the T=4 could elicit some antibodies that are not recognized by T=3 virions. Ultimately, when a patient is immunized with T=4 VLPs, the immune response with virions could effectively be lower.

## ACKNOWLEDGEMENTS

We acknowledge the excellence cluster CellNetworks (Cryo-EM network) of the University of Heidelberg for cryo-EM data collection, the EM core facility at DKFZ, and Baden-Württemberg High Performance Cluster (bwHPC). We thank David Bhella for assistance with structural refinements; Anna Koromyslova for EM images of GII.4 virions; and Benedikt Wimmer for setting up the cryo-EM software. The funding for this study was provided by the CHS foundation; the Baden-Württemberg Stiftung (GLYCAN-BASED ANTIVIRAL AGENTS); Deutsche Forschungsgemeinschaft (DFG, FOR2327); and the BMBF VIP+ (Federal Ministry of Education and Research) (NATION, 03VP00912).

